# Forage preference in two geographically co-occurring fungus gardening ants: a dietary DNA approach

**DOI:** 10.1101/2024.12.17.628420

**Authors:** Matthew Richards-Perhatch, Elizabeth Boshers, Matthew Greenwold, Jon N. Seal

## Abstract

Traditional methods of forage identification are impractical with non-leafcutting fungus gardening ants, making diet-related ecological and life history questions difficult to study. To address this limitation, we utilized dietary DNA metabarcoding on excavated ant fungus gardens to generate forage diversity metrics for the two co-occurring species *Trachymyrmex septentrionalis* and *Mycetomoellerius turrifex*. Ten fungus garden samples from each species were collected from a 60x70 m plot in East Texas. Each of the colonies we sampled was paired with a colony from the other species within 3 m of it. Plant forage diversity was assessed with chloroplast *trnL* primers, and insect frass forage diversity was assessed with mitochondria *COI* primers. DNA metabarcoding identified a total of 44 plant taxa across all samples, but performed poorly when characterizing foraged insect frass. Plant beta diversity was significantly different between the gardens of *T. septentrionalis* and *M. turrifex* colonies, as well as paired colonies. Colony pairs also had significantly different plant alpha diversity. This indicates that diet preference is likely driven both by ant species-specific plant preference, and colony location-specific plant resource availability. Overall, our results show that dietary DNA techniques are a promising tool for the identification of plant forage in ant fungus gardens, enabling the study of future diet-based ecological and natural history questions.

## Introduction

### Fungus Gardening Ants

Fungus-gardening ants are a New World monophyletic group characterized by their cultivation, and obligate consumption, of fungi. These ants use material foraged by their colony as a substrate to grow their fungus garden, feeding off the fungus instead of directly consuming the forage themselves. The more basal fungus-gardening ants forage insect-feces, dry plant debris, leaf-litter, and seed fragments [1,2]. More derived fungus gardening ants practice “higher agriculture” and can be subdivided into two main groups: the leafcutting ants, and the non-leafcutting higher agriculture ants [3,4]. The most specialized of these two groups, the leafcutting ants, are known to forage extensively on freshly cut vegetative material; while the non-leafcutting higher agriculture ants often forage considerable amounts of freshly fallen flower petals, soft leaf tissue, and fruit, in addition to material similar to that collected by more basal fungus gardening species [2,5].

### Traditional Methods of Diet Identification

Visual identification of material carried by foraging workers is a commonly utilized technique in determining ant diet and forage preferences. However, reliance on this technique presents several issues. Visual forage identification is time consuming, relies heavily on taxonomic expertise, and requires foragers to be conspicuous enough for observation [6,7]. In addition, observing individual foragers only offers a small snapshot of total colony dietary breadth, as a single colony may send out hundreds or even thousands of workers to forage [8].

Among fungus gardening ants, visual identification of forage material is most practical with the leafcutting genera *Atta* and *Acromyrmex*, as their foraging strategies predispose them to easy observation. The characteristic foraging trails created by *Atta* and *Acromyrmex* colonies can often be cleanly followed from the vegetation they are cutting to their nest/tunnel entrances, allowing for species level identification of their forage material [9–11]. Unfortunately, visual identification of forage material falls short with fungus gardening ant species outside of *Atta* and *Acromyrmex*. Most non-leafcutting fungus gardening ants do not create true foraging trails, making it difficult and very time consuming to accurately trace the species of vegetation that they are foraging from [12]. The foragers of non-leafcutting fungus gardening ants are also usually smaller and more cryptic than those of leafcutting ants. Additionally, some foragers may collect items that are unsuitable, if not toxic, and deposit them in their nests but not the fungus gardens [41]. Thus observations on forager behavior alone are not likely to depict a comprehensive overview of what ants are collecting and incorporating into their fungus garden. These factors have contributed to a deficit of literature regarding the diet composition of non-leafcutting fungus gardening ants, with the little information available primarily consisting of a handful of forage material observations, often originally published in conjunction with the original natural history descriptions of the ants [1,2]. Clearly, relying solely on visual identification of forage material is insufficient for the study of non-leafcutting fungus gardening ants.

### Dietary DNA

Dietary DNA (dDNA) techniques present a possible alternative to visual forage identification. Metabarcoding techniques based on dDNA, DNA extracted from an organism’s gut, stomach, or fecal matter, have emerged as an accurate and resource-effective way to determine animal diets [7,13]. Previous studies, in multiple animal groups, have successfully utilized dDNA analysis to determine diet composition when direct observations of feeding behavior and/or morphological identification of forage material was insufficient [7,13].

Dietary DNA analysis has only been used to examine ant diet composition in a handful of studies, with even fewer of these studies having attempted to determine broad diet composition [8,14,15]. One factor complicating dDNA ant research is a yet unidentified compound in the gaster of adult ants that has been found to inhibit PCR, potentially impacting gut content dDNA analysis in adult ants [15,16]. As an additional complication, in many species adult ants are unable to digest solid food, instead feeding it to their brood. The adult ants then consume the digested liquid nutrients via trophallaxis from their larvae [8,17]. Wulff et al. (2021) leveraged this adult-larvae digestion dynamic to their advantage when using dDNA analysis to investigate the diet composition of *Solenopsis invicta* [8]. They did so by sequencing the gut content of *S.invicta* larvae instead of adults, as the larvae act as a centralized depository of colony prey and are not known to produce PCR inhibitors [8,15,16].

A similarly innovative approach could be used to analyze the diet composition of fungus gardening ants, since their fungus gardens also act as a centralized depository for colony forage. The fungus gardens cultivated by fungus gardening ants function as an external ancillary digestive system for the colony [2,17–19]. Within this ancillary digestive system, forage materials used as fungal substrate become highly degraded, due both to physical mastication by the ants and enzymatic breakdown by the fungus. This degradation necessitates the use of specialized genetic techniques, like dDNA metabarcoding, that are designed to work with highly degraded staples [2,5,20].

### Evaluation of Diet and Study Species

Using dDNA metabarcoding techniques to identify fungal substrate in ant gardens could be used to answer a number of ecological and natural history questions. Dietary DNA metabarcoding has been successfully used to detect niche partitioning, and overlap, in several animal species [21–25]. These techniques have also been used to understand seasonal and/or biogeographic based shifts in animal diets [26–32].

Dietary differences have been investigated (without the use of dDNA methods) in several species of fungus gardening leafcutting ants. In a dry tropical forest in Costa Rica, the co-occurring leafcutter species *Atta cephalotes* and *Acromyrmex octospinosus* were found to forage almost entirely different resources, with the only exception being they both foraged flowers [33]. A similar trend was observed in Panama, with *Atta* species foraging primarily on tree leaves and co-occurring *Acromyrmex* species foraging primarily on flowers and herbal leaves [34]. To our knowledge, no previous studies have examined differences in diet between non-leafcutting fungus gardening ants.

*Trachymyrmex septentrionalis* and *Mycetomoellerius turrifex* are two higher agriculture non-leafcutting fungus gardening ant species, with a partial overlap of their geographic distribution occurring in East Texas [35,36]. The range of *T. septentrionalis* stretches from Texas to Florida, and as far north as central Illinois and New York; while the range of *M. turrifex* extends from Texas to northeastern Mexico and western Louisiana [36–38]. Like most non-leafcutting fungus gardeners, specific information on the preferred forage material of *T. septentrionalis* and *M. turrifex* is highly limited [2]. What documentation does exist indicates that they both forage insect frass and plant material, but only *T. septentrionalis* has been observed cutting/foraging fresh vegetation [2,39–42]. Their geographic range overlap and the similarity of their documented forage material makes these two species excellent subjects to test the efficacy of using dDNA analysis to study diet in non-leafcutting fungus gardening ants.

The primary objective of this study was to explore whether dDNA metabarcoding could be used to identify and describe the diversity of plant material and/or insect frass used as substrate in the fungus gardens of co-occuring *T. septentrionalis* and *M. turrifex* colonies. The secondary objective of this study was to evaluate whether diversity metrics based on identified fungal substrate were sufficient to detect differences in diet between *T. septentrionalis* and *M. turrifex* colonies.

## Materials and Methods

### Sample Collection

The fungus gardens of ten *T. septentrionalis* and ten *M. turrifex* colonies were sampled from an approximately 60x70 m plot on the University of Texas at Tyler campus in Tyler, Texas, USA.

Sampling occurred from May 16-26, 2023. Each of the ten *M. turrifex* colonies we sampled was paired with a *T. septentrionalis* colony within 3 m of it. These colony pairs, composed of one *M. turrifex* colony and one nearby *T. septentrionalis* colony, were sampled concurrently. Each colony was only sampled once.

Fungus gardens were non-destructively sampled from each colony by careful excavation of the topmost chamber containing an appreciable amount of fungus. Sampling was focused on chambers closer to the surface since deeper chambers were unlikely to contain fungus until later in the season when the upper chambers are too hot for fungus garden growth. Sterilized tools were used when removing fungus gardens from the chamber to minimize DNA contamination from environmental sources and previously sampled colonies. Excavated fungus garden samples were placed in sterile vials containing 95% ethanol and stored in a -80°C freezer.

### Sample Metabarcoding

Metabarcoding was conducted by Jonah Ventures (https://jonahventures.com/) using chloroplast *trnL* (forward primer c: 5’ CGAAATCGGTAGACGCTACG 3’, reverse primer h: 5’ CCATTGAGTCTCTGCACCTATC 3’) and mitochondrial cytochrome c oxidase subunit I (*COI*) (forward primer ZBJ-ArtF1c: 5’ AGATATTGGAACWTTATATTTTATTTTTGG 3’, reverse primer ZBJ-ArtR2c: 5’ WACTAATCAATTWCCAAATCCTCC 3’) primers [43,44]. Individual fungus garden samples were shipped overnight in an ice box to Jonah Ventures in Boulder, CO.

At Jonah Ventures, frozen samples were thawed for 1-2 hours before processing. Sample barcodes were recorded and assigned a well within the 96 well plate or numbered extraction tube. Under a laminar flow hood, sterile cotton swabs (Fisher, Catalog No. 22-363-173) were coated with sediment matter, and the swabs were placed in the corresponding extraction plate or tube. Sterile tweezers and pliers were used to handle cotton swabs and remove the wooden ends of the cotton swab before extraction. Plates or tubes were immediately processed or stored in -20°C until the extraction process could be performed.

Genomic DNA from each fungus garden sample was extracted using the DNeasy 96 PowerSoil Pro Kit (384) (Catalog No. 47017) according to the manufacturer’s protocol. Genomic DNA was eluted into 100 µl and frozen at -20°C.

For *trnL* genetic analysis, a portion of the chloroplast *trnL* intron was PCR amplified from each genomic DNA sample using the c and h trnL primers. Both forward and reverse primers also contained a 5’ adaptor sequence to allow for subsequent indexing and Illumina sequencing.

Each 25 µL PCR reaction was mixed according to the Promega PCR Master Mix specifications (Promega, Catalog No. M5133, Madison, WI) which included 0.4 µM of each primer and 1 µl of gDNA. DNA was PCR amplified using the following conditions: initial denaturation at 94°C for 3 minutes, followed by 40 cycles of 30 seconds at 94°C, 30 seconds at 55°C, and 1 minute at 72°C, and a final elongation at 72°C for 10 minutes.

For *COI* genetic analysis, a portion of the *COI* gene was PCR amplified from each genomic DNA sample using the ZBJ-ArtF1c and ZBJ-ArtR2c primers. Both forward and reverse primers also contained a 5’ adaptor sequence to allow for subsequent indexing and Illumina sequencing. Each 25 µL PCR reaction was mixed according to the Promega PCR Master Mix specifications (Promega, Catalog No. M5133, Madison, WI) which included 12.5 µl Master Mix, 1.0 µM of each primer, 3.0 µl of gDNA, and 7.5 µl DNase/RNase-free H2O. DNA was PCR amplified using the following conditions: initial denaturation at 94°C for 5 minutes, followed by 45 cycles of 30 s at 94°C, 45 s at 45°C, and 45 s at 72°C, and a final elongation at 72°C for 10 minutes.

To determine amplicon size and PCR efficiency, each reaction was visually inspected using a 2% agarose gel with 5 µl of each sample as input. Amplicons were then cleaned by incubating amplicons with Exo1/SAP for 30 minutes at 37°C followed by inactivation at 95°C for 5 minutes and stored at -20°C.

A second round of PCR was performed to complete the sequencing library construct, appending with the final Illumina sequencing adapters and integrating a custom sample-specific 12-nucleotide index sequence (one index read per amplicon) generated by Jonah Ventures. The indexing PCR included Promega Master mix, 0.5 µM of each primer and 2 µl of template DNA (cleaned amplicon from the first PCR reaction) and consisted of an initial denaturation of 95°C for 3 minutes followed by 8 cycles of 95°C for 30 seconds, 55°C for 30 seconds and 72°C for 30 seconds.

Final indexed amplicons from each sample were cleaned and normalized using SequalPrep Normalization Plates (Life Technologies, Carlsbad, CA). 25 µl of PCR amplicon was purified and normalized using the Life Technologies SequalPrep Normalization kit (Catalog No. A10510-01) according to the manufacturer’s protocol. Samples were then pooled together by adding 5µl of each normalized sample to the pool.

Sample library pools were sent for sequencing on an Illumina MiSeq (San Diego, CA) at the Texas A&M Agrilife Genomics and Bioinformatics Sequencing Core facility using the v2 500-cycle kit (Catalog No. MS-102-2003). A PhiX concentration of 15% was used. As a quality control measure, a bioanalyzer chip was used for concentration adjustment at the sequencing center prior to sequencing.

Raw sequence data were demultiplexed using pheniqs v2.1.0, enforcing strict matching of sample barcode indices (i.e, no errors) [45].

### Sequence Processing

After receiving the demultiplexed data in the form of paired end FASTQ files from Jonah Ventures, the files were imported into Qiime2 for further processing (Figure 1) [46].

**Fig 1.**
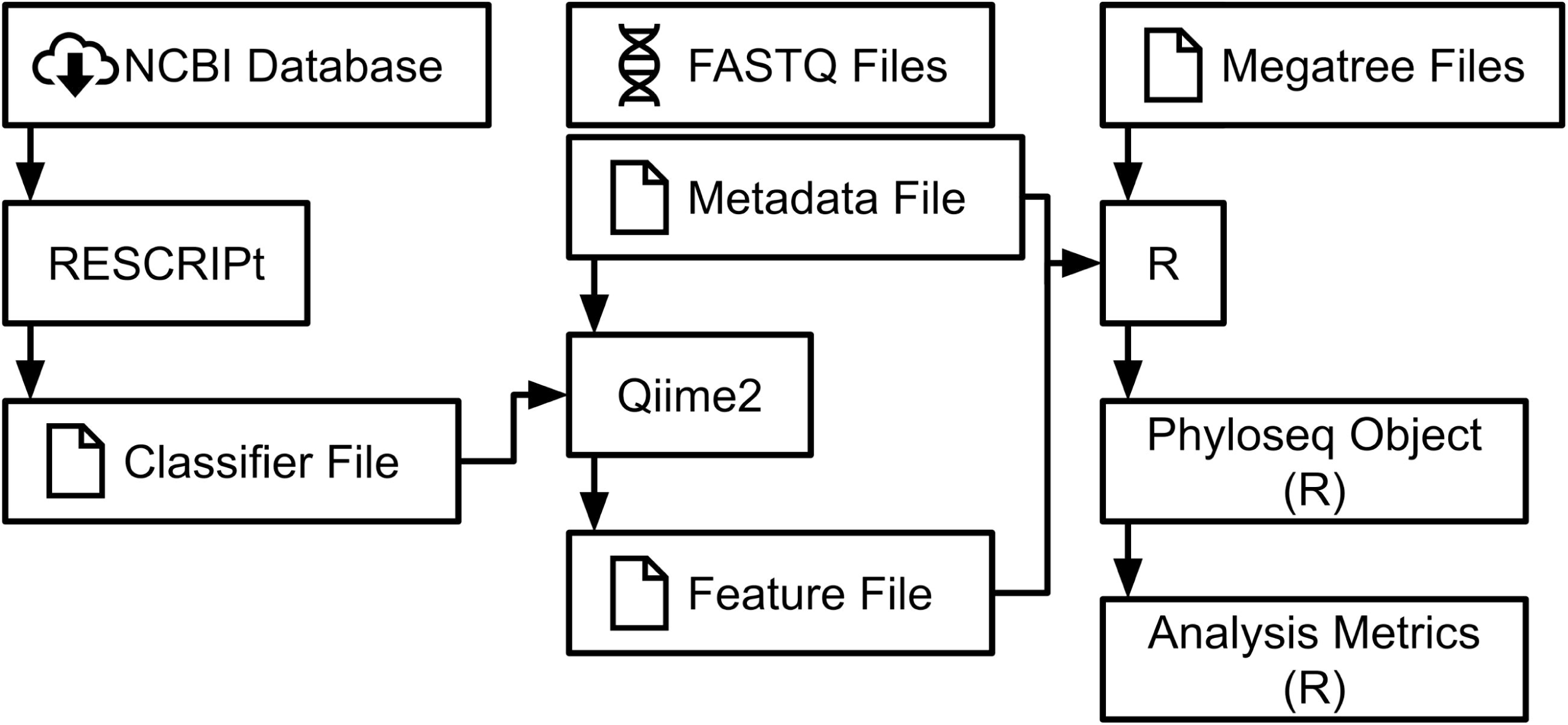
Flowchart of each major bioinformatic step in the development of our analysis metrics.

The Qiime2 plugin “cutadapt” was used to remove primers from the sequences. The Qiime2 plugin “dada2” was used to denoise the *trnL* and *COI* sequences, truncate the *COI* sequences, and merge the forward and reverse *COI* sequence pairs [47]. Only the forward *trnL* sequences were used because of the short *trnL* amplicon length (143 bp). The *trnL* sequences were not truncated because their untruncated median quality scores were 30 or greater. The *COI* sequences were truncated where their median quality scores dropped below 30 (forward: 100 bp, reverse: 87 bp). Rarefaction of the sequences was conducted using Qiime2’s ‘feature-table rarefy’ command. A sampling depth of 9677 was used for *trnL* sequences (Figure 2). Due to low *COI* sequencing depth (<1000) in the majority of samples, downstream analysis was not conducted for *COI* outside of taxonomic classification (Figure 3).

**Fig 2.**
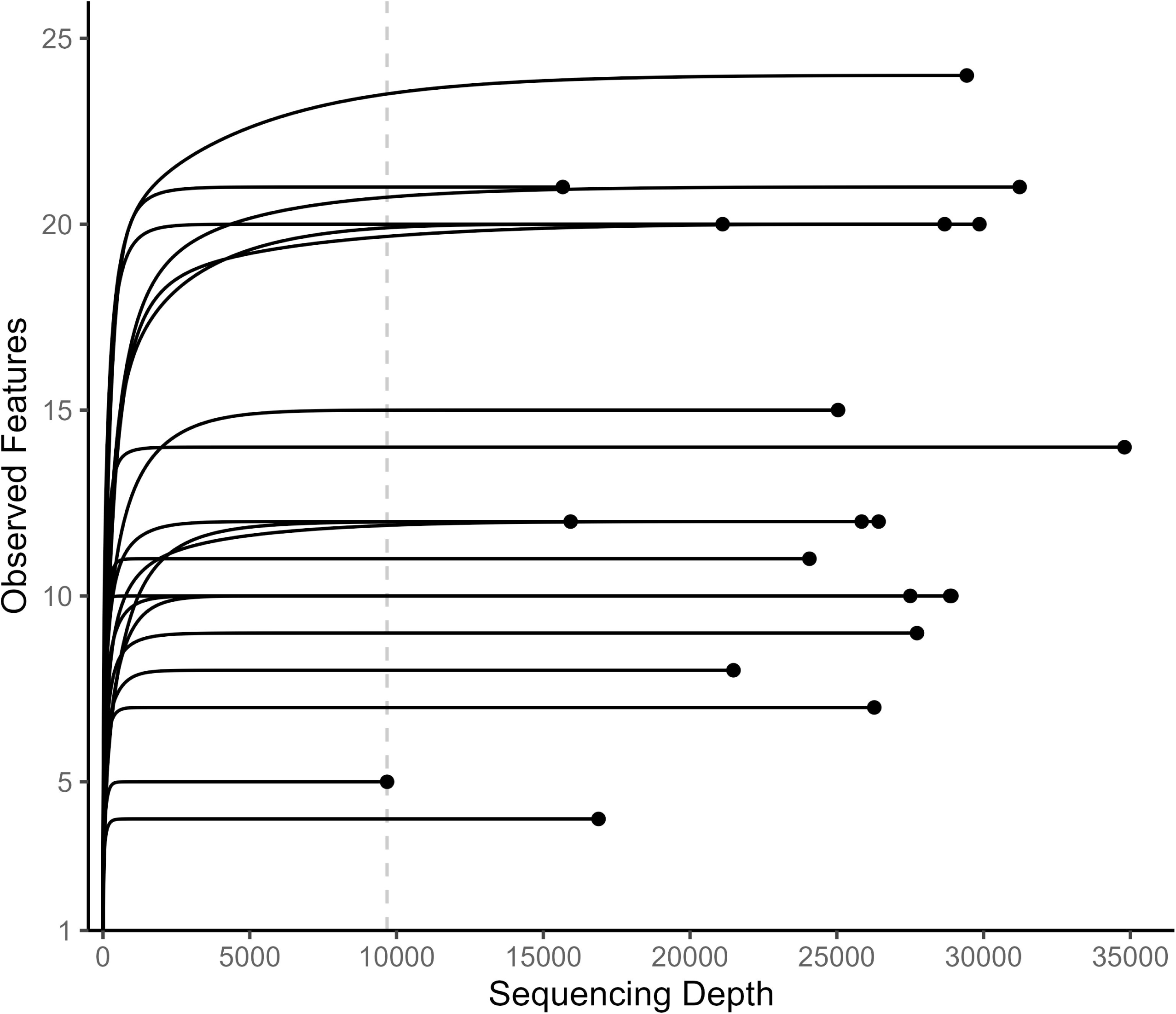
Rarefaction curves for *trnL* sequences. The dashed vertical line shows the chosen *trnL* rarefaction depth of 9677.

**Fig 3.**
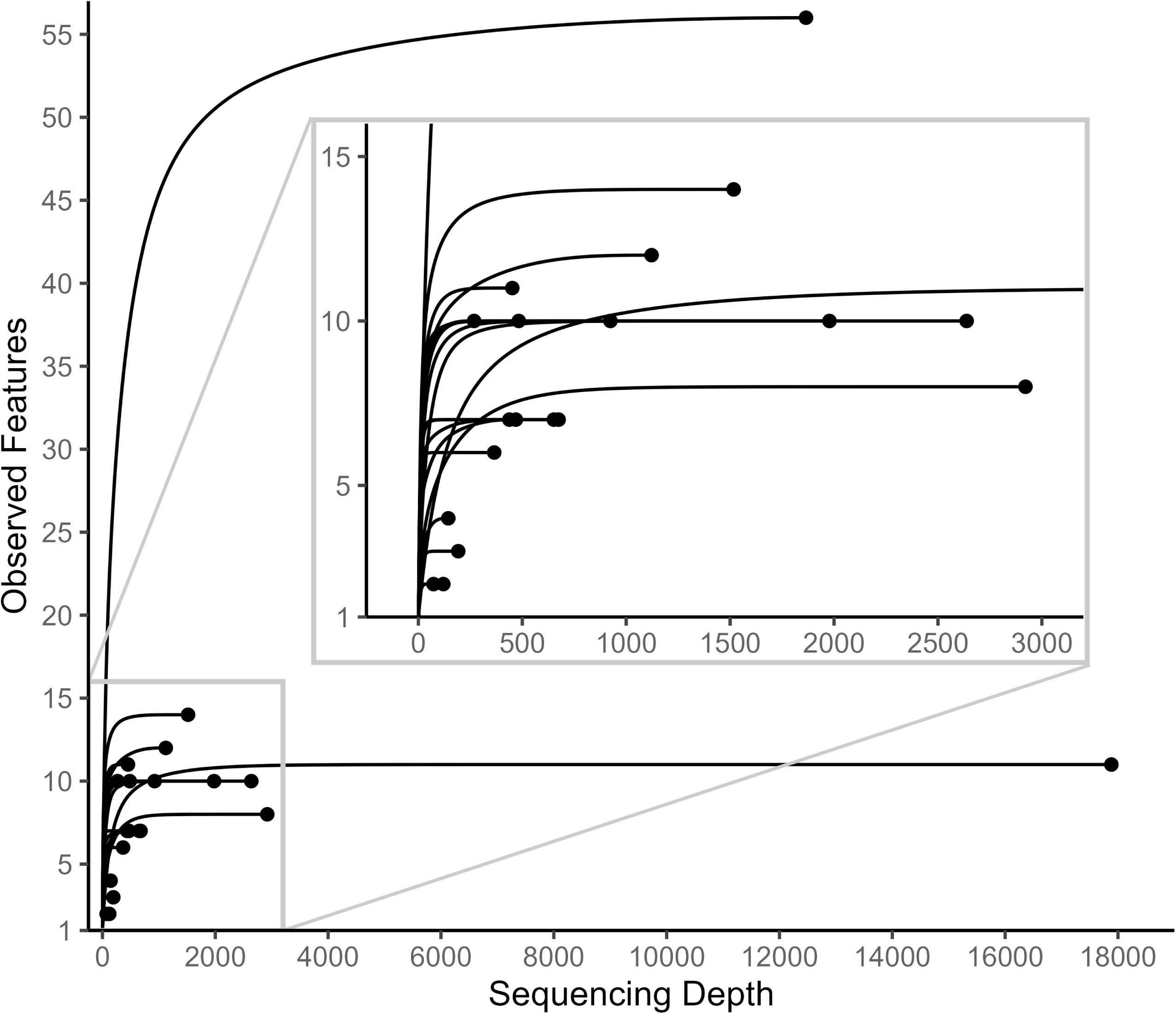
Rarefaction curves for *COI* sequences. The lower left corner of the plot has been magnified to show greater detail.

Qiime2’s “RESCRIPt” plugin was used to create two reference databases from NCBI; one specific to *trnL* and a second specific to arthropod *COI* [48,49]. Initial reference pools were constructed from these downloaded databases via primer-pair search using Qiime2’s ‘feature-classifier extract-reads’ command. These initial pools were both expanded twice using RESCRIPt’s ‘extract-seq-segments’ command. During reference pool expansions of the *trnL* specific database a ‘--p-perc-identity’ value of 0.7 was used. During reference pool expansions of the *COI* specific database a ‘--p-perc-identity’ value of 0.9 was used. These ‘--p-perc-identity’ values were chosen because at the end of their respective reference pool expansions they had extracted sequence segments from ∼70% of their original query sequences [49]. Dereplication during each of these steps was performed using the ‘--p-mode super’ modifier in RESCRIPt’s ‘dereplicate’ command. These expanded reference pools were then used to construct two classifiers, one for *trnL* and one for *COI*, using RESCRIPt’s ‘evaluate-fit-classifier’ command (Figure S1).

Taxonomic assignment for *trnL* and *COI* sequences was conducted using the custom RESCRIPt classifiers described above, in combination with Qiime2’s ‘feature-classifier classify-sklearn’ command.

### Statistical Analysis

Once taxonomic assignment was completed, the Qiime2 feature and metadata files were imported into R and used to create two phyloseq objects using the R packages “qiime2R” and “phyloseq” [50–52]. An independent phylogenetic tree was constructed by grafting our classified *trnL* sequences onto the GBOTB.extended.WP.tre megatree, reported by Jin and Qian (2022), using the ‘phylo.maker’ command from the R package “U.PhyloMaker” [53,54]. Six classified *trnL* sequences failed to graft to the tree due to low taxonomic resolution and were discarded.

The resulting phylogenetic tree was converted to a phylo class object using the ‘phyloseq::read_tree’ command; multichotomies were then resolved using the ‘multi2di’ command from the R package “ape” [55]. This tree was merged with our phyloseq object using the ‘phyloseq::merge_phyloseq’ command. This updated phyloseq object was used to calculate alpha diversity for each colony, in the form of Faith’s phylogenetic diversity (Faith’s PD) scores and species richness, using the ‘calculatePD’ command in the R package “biomeUtils” [56].

Beta diversity between each colony was also calculated from the updated phlyoseq object, in the form of unweighted unique fraction metric (uUniFrac) dissimilarity distances, using the ‘phyloseq::distance’ command.

These specific diversity metrics were chosen for two primary reasons: 1) their calculations are based on relative frequency of occurrence (binary presence/absence) of the sequenced taxa, and 2) their calculations take into account the relative relatedness of the sequenced taxa.

The number of sequencing reads from dDNA often does not accurately represent the biomass of consumed material [6,13]. This is particularly relevant in studies like ours, where sequenced material consists of a broad diversity of partially digested multi-cellular samples. It is likely that digestive processes, the order items are consumed, primer biases, and taxonomic biases in the reference database influence both the sequencing outcomes and taxonomic classification [6,57,58]. Due to the known impact of these factors on estimations of abundance, only metrics based on binary presence/absence of the sequenced taxa (i.e. uUniFrac and Faith’s PD) were used.

Another issue arises from the various states of DNA degradation present in our samples; as sequences originating from the same species, or even the same individual, may end up classified at different taxonomic levels. For example, if one group of sequences are classified as *Fagaceae* and another group of sequences from the same sample are classified as *Fagaceae*

*Quercus*, it is not currently possible to determine if this indicates the presence of two separate genera of *Fagaceae*, or if both sets of sequences come from the same genera. This can result in the artificial inflation of identified taxa, as sequences from a single taxon could be functionally double counted. Some previous dDNA studies have attempted to account for this issue by compressing their post-classified dDNA taxa into diet events: instances where there is unequivocal evidence from dDNA analysis that an organism consumed at least one individual of a species [8,59]. While this approach does minimize double counting of taxa, it increases the risk of undercounting dietary events and can obscure dietary events of taxa unable to be identified to the species level [59]. To avoid the drawbacks of using a diet event based approach, we instead utilized metrics that included the phylogenetic relationship of classified taxa in their calculations (i.e. uUniFrac and Faith’s PD). Using these phylogeny-weighted metrics allows sequences classified at different taxonomic levels to be assessed as separate taxa, while minimizing the impact of possible double counts.

Our analysis incorporated two main predictor variables: colony ant species (*T. septentrionalis* and *M. turrifex*) and colony pair ID (Pair-2, Pair-8, Pair-12, Pair-14, Pair-17, Pair-20, Pair-28, Pair-30, Pair-44, and Pair-50). We tested for differences in fungal substrate plant taxa alpha diversity (Faith’s PD and species richness) and beta diversity (uUniFrac dissimilarity distance).

Faith’s PD and species richness distributions were tested for normality via the Shapiro-Wilk normality tests by using the ‘shapiro.test’ command in R. To analyze Faith’s PD and species richness against each of our predictor variables, several analysis of variance (ANOVA) analyses were conducted using the ‘aov’ command in R.

Permutational analysis of variance (PerMANOVA) analyses were conducted using the ‘adonis2’ commands in the R package “vegan” [60]. Our PerMANOVA analyses used 9,999 permutations.

Data visualization was performed with the R packages “tidyverse”, “ggtree”, and “ggrepel” [61–63].

## Results

### Plant Taxa Identified from Fungus Garden Samples

Across all sampled fungus gardens, a total of 44 plant classifications were identified from *trnL* sequences (Figure 4). Fungal substrate plant taxa composition was derived from 32 plant classifications for *T. septentrionalis* and 33 plant classifications for *M. turrifex*. There was considerable overlap in plant taxa composition between the two species, with 21 plant classifications present in both *T. septentrionalis* and *M. turrifex* colonies. 11 plant classifications were identified only from *T. septentrionalis* colonies and 12 plant classifications were identified only from *M. turrifex* colonies.

**Fig 4.**
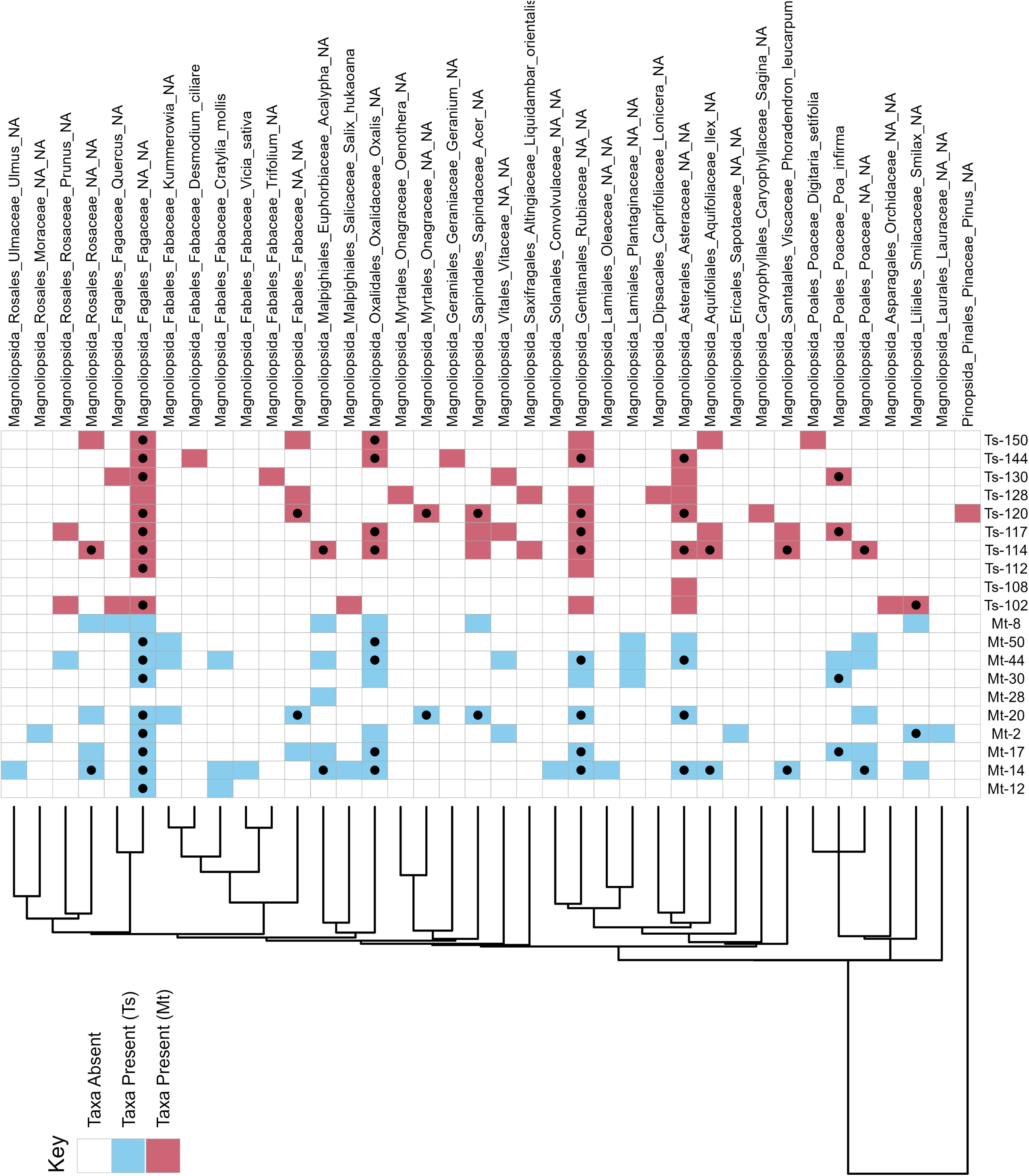
Phylogenetic tree and binary heat map of plant taxa identified from fungus garden samples. Each branch end is linked to a row in the binary heat map, showing presence/absence of each respective plant classification identified in the *M. turrifex* (Mt) and *T. septentrionalis* (Ts) fungus garden samples. Cells with a black dot indicate the plant classification was present in both colonies from that pair.

In *T. septentrionalis* colonies, the most frequently occurring plant classifications (occurred in ≥50% of *T. septentrionalis* samples) were: *Fagaceae*, *Magnoliopsida*, *Rubiaceae*, *Asteraceae*, and *Fagales.* For *M. turrifex* colonies, the most frequently occurring plant classifications (occurred in ≥50% of *M*. *turrifex* samples) were: *Fagaceae*, *Oxalis* (*Oxalidaceae*), *Magnoliopsida*, *Rubiaceae*, *Fagales*, *Acalypha* (*Euphorbiaceae*), and *Poaceae*.

### Plant Taxa Alpha Diversity

ANOVA analyses were used to determine if Faith’s PD or species richness of fungal substrate plant taxa differed significantly between colonies based on colony ant species or colony pair ID. *Trachymyrmex septentrionalis* and *M. turrifex* fungus gardens did not have significantly different Faith’s PD (F_1,18_ = 0.045, p = 0.834), nor did they have significantly different species richness (F_1,18_ = 0.223, p = 0.643). There were, however, significant differences in Faith’s PD (F_9,10_ = 3.921, p = 0.022) and species richness (F_9,10_ = 3.105, p = 0.046) between colony pairs.

### Plant Taxa Beta Diversity

A PerMANOVA analysis was used to determine if the uUniFrac dissimilarity distances of fungal substrate plant taxa were significantly different between colonies based on colony ant species or colony pair ID. There was a significant difference found between colony ant species and the uUniFrac dissimilarity distances of their plant taxa (p = 0.046) (Figure 5). There was also a significant difference between colony pairs and the uUniFrac dissimilarity distances of their plant taxa (p = 0.014).

**Fig 5.**
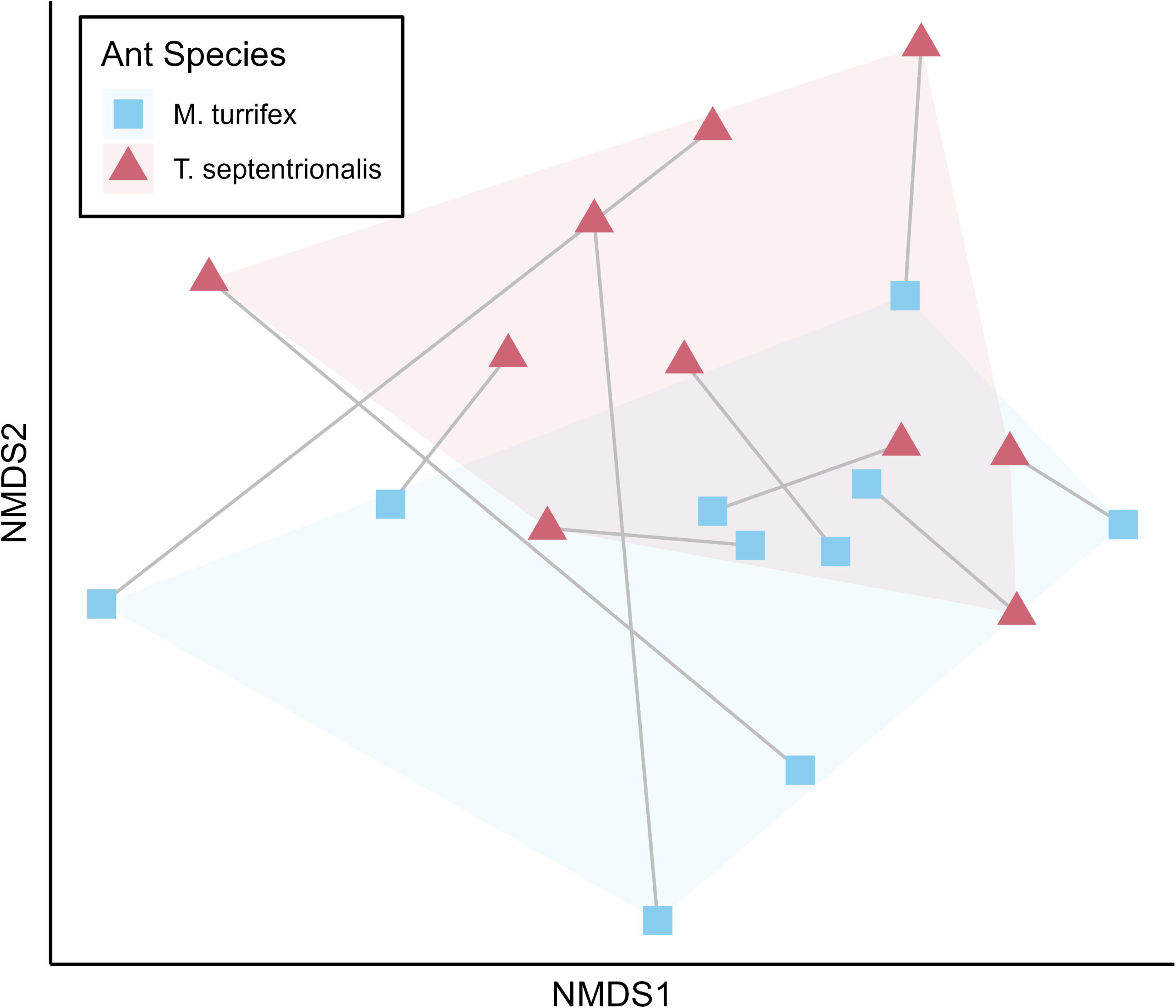
NMDS plot showing the uUniFrac dissimilarity of plant taxa identified from fungus garden samples. Colony pairs are connected by grey lines.

### Arthropod Taxa Identified from Fungus Garden Samples

A total of six arthropod classifications were identified from *COI* sequences. All colony fungus garden samples contained *Insecta* sequences. Only four samples contained sequences identified to a lower taxonomic level than *Insecta*: *M*. *turrifex* colony 44’s fungus garden contained *Carabidae* (*Coleoptera*) and *Lepidoptera* sequences, *M*. *turrifex* colony 14’s fungus garden contained *Anomoea laticlavia* (*Coleoptera Chrysomelidae*) sequences, *M*. *turrifex* colony 20’s fungus garden contained *Coleoptera* sequences, and *T. septentrionalis* colony 112’s fungus garden contained *Hemiptera* sequences.

## Discussion

Dietary DNA techniques successfully identified plant and arthropod taxa present in the fungus gardens of *T. septentrionalis* and *M. turrifex*. The dDNA analysis was particularly successful with the *trnL* primer, as it identified a relatively large number of plant taxa, many of them to a relatively high level of taxonomic resolution. These results could be particularly impactful considering the literature deficit of specific plant taxa known to be foraged by non-leafcutting fungus gardening ants. Only a handful of previous studies make any mention of specific foraged plant taxa, with most only describing the general type of material the ants were collecting such as flower petals or dead leaves [1,2,40–42,64–68].

Colony ant species appeared to be a significant factor in beta diversity metrics, but not alpha diversity metrics, of fungal substrate plant taxa. Similarly, Colony pair appeared to be a significant factor in both plant taxa beta diversity metrics and alpha diversity metrics. These results suggest that diet preference is likely driven both by ant species-specific plant preference and colony location-specific plant resource availability. The presence of dietary differences between *T. septentrionalis* and *M. turrifex* colonies on our site appears similar to short-term observations of dietary differences made on co-occurring *Atta* and *Acromyrmex* colonies [33,34].

The plant taxa we identified from *T. septentrionalis* and *M. turrifex* fungus garden sequences were consistent with the limited existing descriptions of forage material collected by these ant species. One previous study observed *M. turrifex* collecting *Sorghum halepense* and *Solidago sp.*, both of whose families, *Poaceae* and *Asteraceae* respectively, were present in our samples [40]. Another previous study suggested that *T. septentrionalis* collects *Quercus* (*Fagaceae*), which was also present in our samples [41].

If we include descriptions of known forage material from other higher agriculture non-leafcutting ants, we find additional overlap between our identified plant taxa and that of previous studies. *Lauraceae*, *Rosaceae*, *Rubiaceae*, *Poaceae*, and *Euphorbiaceae* are all plant families which were identified in our samples, and are known to be collected by or, in the case of lab colonies, accepted by higher agriculture non-leafcutting ants [64,66]. Leafcutting species like *Atta texana* are known to forage on *Pinus* (*Pinaceae*) needles, but our study is the first published record of a non-leafcutting species collecting *Pinus* [69]. The ants may be collecting pieces of male pine cones, as this has been observed in the field and laboratory during early spring (March-April) (JN Seal, unpublished observations).

A notable drawback of only using dDNA analysis on sampled fungus gardens is that dDNA analysis alone cannot determine the state plant material was in when it was collected by the ants. Some higher agriculture fungus gardeners outside of *Atta* and *Acromyrmex*, like *T. septentrionalis*, are known to cut fresh vegetative material to use as fungus garden substrate, while others do not [2,41]. In our study, we could not determine whether the detected plant sequences originated from the ants foraging fresh cut plant material, plant detritus, or frass. We also could not determine which part of the plant they collected (i.e. leaves, flowers, or sticks).

The inability to detect which state plant material was collected in could obscure differences in foraging behavior between these two ant species. For example, one ant species may forage on *Quercus* (*Fagaceae*) catkins, while a second ant species may forage caterpillar frass that contains *Quercus* leaves. Similarly, one ant species may cut soft *Quercus* vegetation, whereas a second ant species might only take dead vegetation that has fallen from the tree. In all of these examples the ant’s fungus gardens would contain *Quercus* sequences, but dDNA analysis would be unable to differentiate the state the material was collected in. Additionally, since the number of sequencing reads cannot be reliably used as a quantitative measure of the biomass of digested material, we cannot use dDNA to determine if colonies are foraging different amounts of the same material [6].

While our study establishes dDNA analysis as a potential tool for identification of fungus garden substrate, we also identified several limitations that should be accounted for in future studies attempting to use similar techniques. One issue was the low taxonomic resolution of some plant sequences. While we were able to successfully identify 44 plant classifications in the fungus gardens using our selected *trnL* primer, unidentified *Magnoliopsida* had the sixth highest absolute read abundance post-rarefaction, indicating that a number of plant species may be present in the fungus gardens that our *trnL* primer was unable to identify to an informative taxonomic level. In future studies this could be mitigated, at least in part, by using multiple plant primers designed for different target regions [23,70,71]. Another issue was the misidentification of plant sequences at the species level. While some identified plant species, like *Desmodium ciliare* (*Fabaceae*), are known to occur on our study site; other species, such as *Salix hukaoana* (*Salicaceae*), are not (though other species of *Salix* are certainly in the general vicinity). These errors were likely the result of our classifier misidentifying sequences as related taxa due to an incomplete reference database. Ideally, future studies could construct a region-specific taxonomic library of plant sequences to be used when training their classifier [72]. Barcoding insect species present in a study site would be more difficult, as they are more mobile and can inhabit above both and below ground areas during different life stages.

Dietary DNA analysis alone is not capable of distinguishing *trnL* sequences from plant material foraged directly by the ants and *trnL* sequences from insect frass foraged by the ants. Since *Trachymyrmex septentrionalis* and *M. turrifex* both readily forage insect frass containing plant material, plant DNA sequences from foraged frass were potentially detected by the *trnL* primer we used [13,40,42]. We had hoped to use the *COI* primer results to gain insight into what arthropod taxa the ants were collecting frass from, but the primer performed too poorly to draw any meaningful conclusions. Other studies attempting to determine *Lepidoptera* species from their frass found that amounts of dry frass less than 5 mg may be too small to detect insect DNA sequences [73]. This insect sequence detectability cutoff was likely higher in our samples due to the physical and enzymatic breakdown of foraged material occurring in the fungus gardens, making it difficult to detect frass-producing-insect DNA [2]. Another potential confounding issue could be a decrease in PCR efficiency due to our use of a 1-step PCR approach versus a 2-step PCR approach for sequencing (See Methods Bohmann et al. (2022) for a discussion on the topic) [74].

Many of the issues we discussed can be mitigated by taking the limitations of dDNA analysis into account when designing future studies. Our recommendations for future studies include: 1) using multiple plant primers to maximize taxonomic coverage, 2) constructing a region-specific taxonomic library of plant sequences, 3) sampling at several different time points to allow detection of temporal changes in fungal substrate plant composition, and 4) incorporating a supplemental observational component to gain an understanding of what state forage material is being collected in. Overall, we have demonstrated the efficacy of using dDNA analysis as a tool for the taxonomic identification of forage material in fungus-gardening ants, and as a possible tool to detect dietary differences between species. Dietary DNA analysis offers an opportunity to improve our understanding of plant taxa foraged by non-leafcutting fungus gardening ants, and, with appropriate supporting methods, can be applied to gain new insight into their ecology and life history.

## Acknowledgments

We would like to thank Christine Bays, Dillon Flowers, Gabriel McDanield, and Grant Rook, all of whom took time out of their lives to help with colony excavation and sampling.

## Data Availability

The raw sequences for all samples have been deposited with links to BioProject accession number PRJNA1124954 in the NCBI BioProject database (https://www.ncbi.nlm.nih.gov/bioproject/PRJNA1124954). The Qiime2, RESCRIPt, and R code used during this project can be found at https://github.com/mrichardsperhatchv/FORAGE-PREFERENCE-IN-TWO-GEOGRAPHICALLY-CO-OCCURRING-FUNGUS-GARDENING-ANTS.

## Supplemental

**Fig S1. The F-measure of our classifiers being used on their own training dataset.**

