## Supplementary figures and images for "Forage preference in two geographically co-occurring fungus gardening ants: a dietary DNA approach"

### Supplemental Figure 1

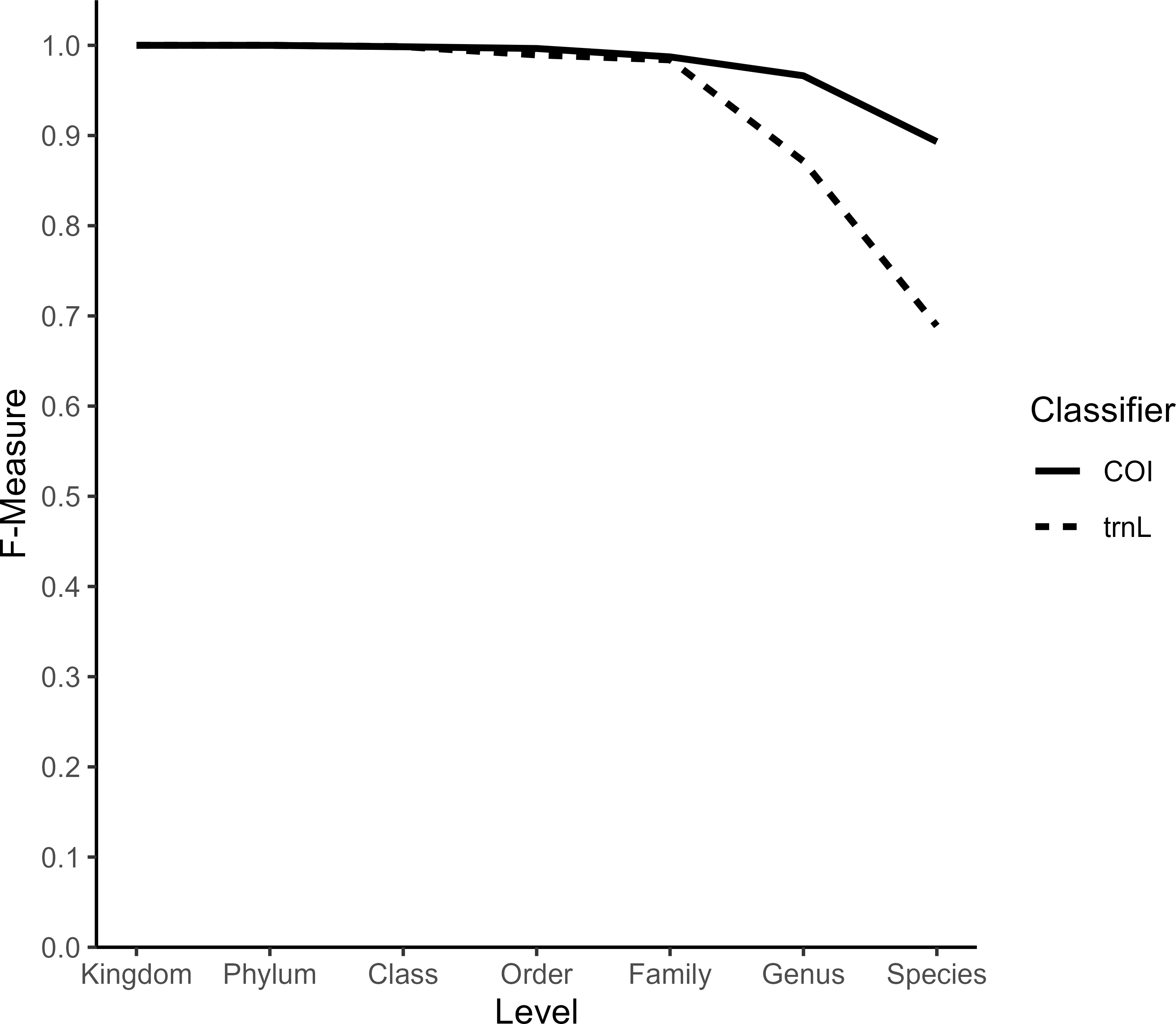
